# Tree diversity effects on forest productivity: disentangling the effects of tree species addition vs. substitution

**DOI:** 10.1101/2021.05.07.443133

**Authors:** Toïgo Maude, Castagneyrol Bastien, Jactel Hervé, Morin Xavier, Meredieu Celine

**Author notes:** Correspondence author*, Toïgo Maude, ^a^CEFE UMR 5175, CNRS – Université de Montpellier – Université Paul-Valéry Montpellier – EPHE–IRD, 1919 Route de Mende, F-34293 Montpellier, France.

## Abstract

1. Mixture effect on stand productivity is usually apprehended through a substitutive approach, whereby productivity in mixed stands is compared to productivity in monocultures, at equivalent stand density. This approach has proved that in many cases mixed stands perform better than monospecific forests, however, we do not yet have a solid theory about species behaviour in the mixture or even guidelines for combining species. The addition of a second tree species to an existing mono-specific stand has received much less consideration. Yet, this approach has the potential to separate the facilitation effect from the complementarity effect.
2. We compared the effect of tree species substitution vs. addition on the productivity of maritime pine and silver birch in a young tree diversity experiment implemented in 2008 in SW France.
3. Substituting pines with birches to create two-species mixtures resulted in an increase of tree productivity at stand level beyond what was expected from monocultures (i.e., overyielding). In contrast, creating mixture through the addition of birches to pine stands had no effect on the maritime pine stand productivity (transgressive mixture effect not significant). This absence of effect is produced by two distinct density-dependence responses at an individual level.
4. Our results allow clarifying the cases in which a mixed stand can be considered as an alternative to a monoculture of a productive species. In particular, the addition of a pioneer and soil low-demanding species during young developmental stages is a possibility to diversify the stand and potentially to increase ecosystem services without altering the productivity of the target species.

## INTRODUCTION

### 1§ CHALLENGES AND DETERMINANTS OF MIXED PLANTATIONS

Despite ample evidence that mixed stands provide more ecosystem services than monospecific forests under various ecological conditions (Baeten et al., 2019), most planted forests are still managed as monocultures. Moving towards ecologically intensive and sustainable forest management requires a sound understanding of the drivers that could improve or hamper the benefits of mixed forests (Felton et al., 2010). Tree species diversity has well documented positive effects on tree productivity (Gamfeldt et al., 2013). Such positive effects are driven by both complementarity and selection effects (Loreau & Hector, 2001). Complementarity mostly refers to niche partitioning processes whereby mixed stands are better able to capture resources than monospecific stands (Jucker et al., 2015), and to facilitation, where one species in the mixture benefit to the others, e.g. via improved resource quality (N-fixing species) acquisition (water uplifting), or protection against herbivores (Caspersen et al., 2018; Kunz et al., 2019). Selection effect refers to situations where a highly productive species recruited in the mixed stand drives positive mixture effect (Fox, 2005; Loreau & Hector, 2001). However, recent studies have highlighted that positive diversity-productivity relationship is strongly context-dependent. For instance, species functional characteristics or stand structure can modify the shape and strength of the diversity-productivity relationship (Forrester, 2014; Grossman et al., 2017). Disentangling drivers of the mixture effect require innovative conceptual framework supported by novel experimental approaches based on stand density, a major component of stand structure that can be controlled by thinning operations.

### 2§ STAND DENSITY A KEY DETERMINANT OF MIXTURE EFFECT ON STAND PRODUCTIVITY

Stand density influences the degree of canopy closure, which in turn participate in the regulation of light transmittance, the interception of water precipitations, belowground competition for water and can modify understory microclimate, understory vegetation, and soil biodiversity (Baeten et al., 2019; Gaudio et al., 2011; Henneron et al., 2017; Ligot et al., 2014; Perot et al., 2017). Stand density is also a major driver of tree-tree competition, being used to calculate several competition indices in forest (Biging & Dobbertin, 1992). Due to the considerable effects of tree density on canopy packing and abiotic factors in forest stands, it is surprising that only a few studies addressing the effect of tree diversity on productivity in temperate forest explicitly questioned the importance of stand density (Forrester, 2014; Jucker et al., 2016). Yet, complementarity among species and intra-specific competition both intensify with stand density. This was documented in mixed stands of late-successional species (Amoroso & Turnblom, 2006; Forrester et al., 2013) of slow-with fast-growing tree species (Condés et al., 2013) and of species with contrasted shade tolerance (del Rio & Sterba, 2009). However, it remains unclear how the mixture effect can possibly be modified by stand density, especially in young plantations of fast-growing tree species.

### 3§ CONTROLLING STAND DENSITY TO COMPARE MONOCULTURES TO MIXED STANDS: OVERYIELDING, THE CLASSICAL INDEX BASED ON SPECIES SUBSTITUTION

Our understanding of the diversity-productivity relationship in mixed forests is further hampered by unresolved methodological issues. The net biodiversity effect generally simply compares the observed productivity of a mixture to a theoretical mixture assembled with the same proportion of trees drawn from the component monocultures (Loreau, 1998; Loreau & Hector, 2001). As such, overyielding can be seen as a measure of changes in stand productivity due to the substitution of a species by others. Estimating the effect of species mixture on productivity through overyielding has several advantages. First, it provides a quantitative estimate of the net biodiversity effect on stand productivity (Tobner et al., 2016). Second, because it compares the productivity of the mixture to the weighted productivity of the component monocultures, it allows addressing whether the mixture performs better than the average of monocultures (*overyielding*) or the most productive monoculture (*transgressive overyielding*).

### 4§ LIMITATIONS LINKED TO SPECIES SUBSTITUTION AND ITS RELATED OVERYIELDING

The use of overyielding estimate has also several shortcomings. First, because it is inherently defined at the stand level, overyielding does not account for species-specific responses to tree diversity and knowing which species benefits or not from the mixture is of primary importance, particularly when it comes to harvest species at different times because of differences in growth patterns. Still the effects of tree diversity may not be symmetrical (del Rio & Sterba, 2009), which is a major concern to understand the functioning of mixed species forests. As a consequence, considering the mixture effect on species productivity and on individual tree productivity is a first step to disentangle the mechanisms underlying the diversity-productivity relationship (Nadrowski et al., 2010). Moreover, from a practical point of view, the conversion of monocultures to mixed stands through species substitution is not without management problems. On the one hand, the silviculture of mixed stands, particularly in cases of intimate mixing, is complicated by the difference in growth rates of the different species and the lack of knowledge about the optimum growing space for trees of each species. On the other hand, wood product processing chains are often specialized in a limited number of species and may not be able to offer a market for substitute species.

### 5§ SPECIES ADDITION AS AN ALTERNATIVE TO SPECIES SUBSTITUTION

An alternative to species substitution is the addition of a new species within an existing stand. A species addition could be less constraining than species substitution by making it possible to keep the same harvesting rate for the target tree species, for example in the case of alternate-row mixing. Second if resource complementarity cannot be distinguished from facilitation through species substitution, any gain of productivity observed through species addition should be the signature of facilitative processes (i.e. comparison of one tree species productivity with vs. without any heterospecific neighbours). So, either species addition or substitution should be considered to design and manage mixed-species forests and dedicated experiments are needed to disentangle their specific effects on productivity.

### 5§ OBJECTIVES AND TESTED HYPOTHESES

Using a long-term tree diversity experiment, we experimentally uncoupled the effect of species addition vs. substitution on forest stand productivity while controlling for stand and species-specific density to gain further insight into the mechanisms underlying the effect of species addition and substitution. We focused on two-species mixtures of maritime pine (*Pinus pinaster Ait.*) and silver birch (*Betula pendula Roth*) at two stand densities. Although the two species studied are fast-growing species, they are nevertheless distinct in terms of growth dynamics and tree sizes. In the case of species substitution, we expected a positive global mixture effect (ME) with a positive specific effect for both pine and birch. By contrast we anticipated a negative transgressive effect (TME) as birch is notably less productive than maritime pine in the local conditions of the experiment. In the case of species addition, we hypothesized opposite patterns of response: a negative ME due to increased competition between trees (due to higher tree density) but a positive TME due to a tree packing effect and a weak competition from silver birch in pine stands. Lastly, we expected that all mixture effects would intensify with stand density.

## METHODS

Both maritime pine and silver birch are light demanding, fast-growing tree species and native to the site. The area of distribution of maritime pine is mainly restricted to Spain, the south-west of France and the north-west of Italy. Maritime pine is a highly drought tolerant species and a major species of production in France yielded exclusively in monoculture. Conversely silver birch is widely distributed across Europe from the Atlantic to eastern Siberia. Silver birch is yielded in the Northern and Eastern Europe and despite the interest shown by these countries, in the Atlantic this tree species is depreciated (Hynynen et al., 2010).

### EXPERIMENTAL DESIGN

The ORPHEE experiment is located 40km south of Bordeaux (44°440 N, 00° 460 W) and belongs to the worldwide Tree Diversity Network (TreeDivNet). The experimental plantation was established in 2008 on a clear cut of former maritime pine stands on a sandy podzol. Stumps were removed and the site was ploughed and fertilized with phosphorus and potassium before planting. In total, 25 600 trees of five native species (Silver birch, *Betula pendula;* pedunculate oak, *Quercus robur;* Pyrenean oak, *Quercus pyrenaica;* holm Oak, *Quercus ilex* and maritime pine, *Pinus pinaster*) were planted within a 12ha area. Eight blocks were established, with 32 plots in every block, corresponding to the 31 possible combinations of one to five species, with an additional replicate of the combination of five species. Each plot contained 10 rows of 10 trees planted 2 m apart, resulting in 100 trees per plot, with a plot area of 400 m^2^. The total initial stand density was therefore 2500 tree per hectare in each plot. Tree species mixtures were established according to a substitutive design, keeping the total tree number (n=100) equal in all plots. Within plots, individual trees from different species were planted in a regular alternate pattern, such that a tree from a given species had at least one neighbour from each of the other species within a 2 m radius. Plots were three meters apart and were randomly distributed within blocks. The entire experimental site was fenced to prevent grazing by mammalian herbivores.

### PLOT SELECTION

The present study analyses growth data collected in 2014 (7 years-old) on the target trees at the center of the plots in order to avoid edge effects (measured planting locations = 36). Importantly, at this time of plot development, oak trees were on average 112 cm high and have a negligible growth in diameter, whereas pines and birches were on average five times taller than oaks (563 and 510 cm high, respectively). As a consequence, oak trees were confounded with the understorey vegetation. By considering oak seedlings as part of the understorey vegetation we therefore focused solely on birch and pine growth. However we do not deny the existence of belowground interactions as the understorey can represent a large part of the fine root biomass in maritime pine stands (Bakker et al., 2006). But the three oak species represent only a few individuals among the 25 species found in the understorey (i.e. the most common are *Molinia caerulae, Ulex minor* and *Pteridium aquilinum* Corcket et al., 2020), which allows us to reasonably assume that the impact of these relatively few oak individuals on the productivity of pine and birch at these developmental stages is negligible. We tested the effect of species addition and substitution on tree and stand volume by selecting monocultures and mixed stands of birch and pine at equal stand density: the “high-density plots” (2500 t/ha) had 100 pines or 100 birches for monoculture plots or a mixture of 50 pines and 50 birches. The “medium-density plots” (1250t/ha) had 50 pines or 50 birches for monoculture plots, or a mixture of 25 pines and 25 birches. We completed the sampling by selecting “low-density plots” as monocultures (625 t/ha) with 25 pines or 25 birches (figure 1). To avoid bias when comparing volume of mixed stands and monocultures we eventually selected plots with less than 15% of dead trees as an optimal balance between the number of plots per treatments and the number of trees per plot (Supporting Information Table S1).

**Figure 1.**
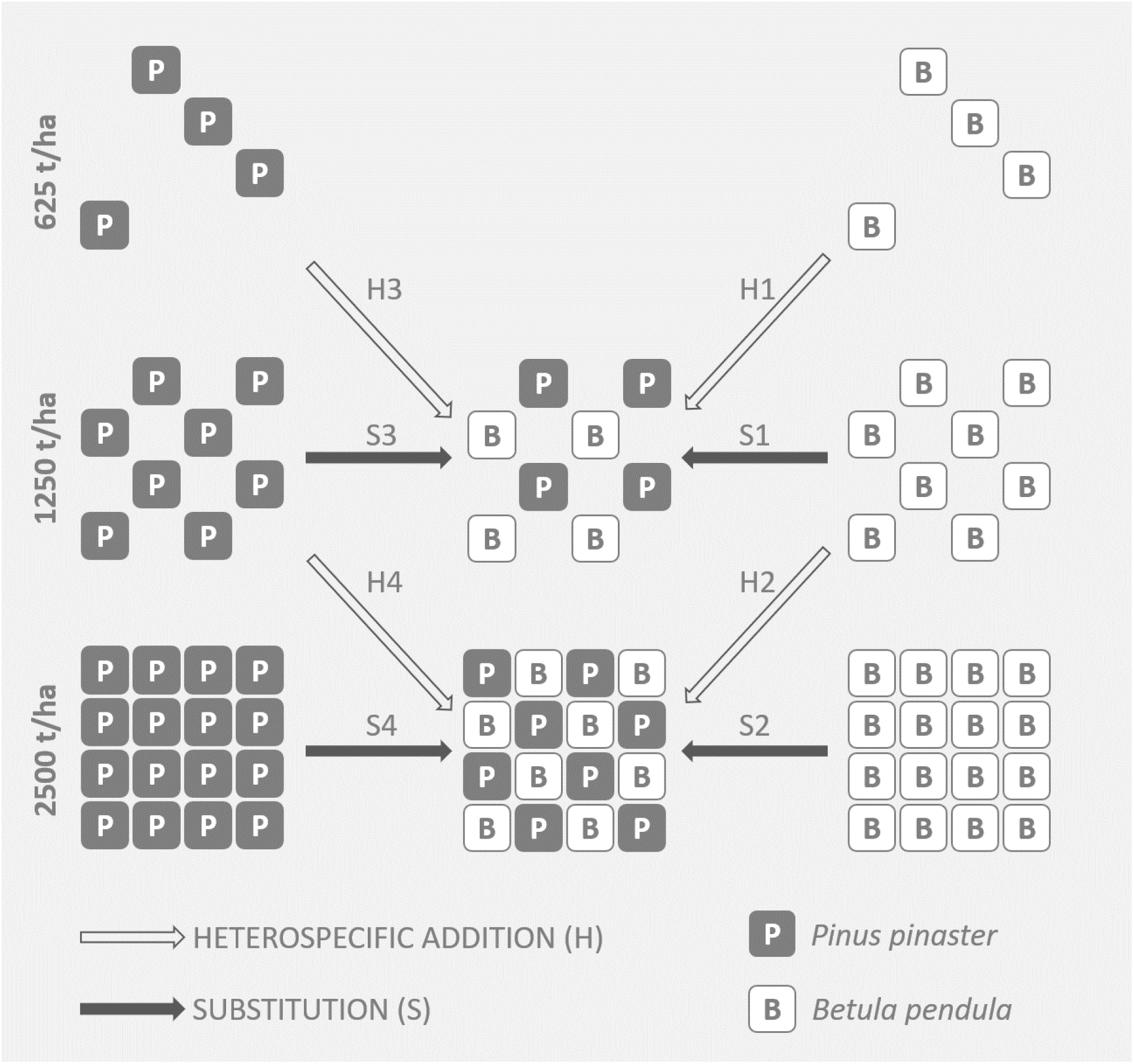
Schematic representation of the experimental treatments consisting in three levels of stand density (low: 625t/ha, medium: 1250t/ha and high: 2500t/ha) and stand compositions, from left to right: *Pinus pinaster* in monocultures, mixed *Betula pentula* - *P. pinaster* stands (proportion of 50% of each species) and monoculture stands of *B. pendula*. Arrows indicate the between-treatment comparisons disentangling heterospecific addition (solid arrows) and species substitution (black arrows). Arrows are numbered according to the different experimental treatments compared in the result section.

### DENDROMETRIC DATA

We measured the height of every 36 innermost planted trees at the center of every plot using a graduate pole, each year from 2008 to 2014. We therefore measured 36, 18 or 9 pines or birches in the high, medium and low-density plots, respectively. We also measured circumferences at 1.30m from 2012 to 2014 on 7 randomly chosen pines and 7 randomly chosen birches per plot, irrespective of plot composition. We used height-circumference relationships to estimate the circumferences of trees that have not been measured in 2014 (Supporting Information Figure S1), then we estimated tree volume following the generic model developed by (Deleuze et al., 2014). We assigned a minimum volume of 0.01m^3^/ha to the few trees below 1.30m in height (corresponding to the minimum volume found in the data set). Eventually we estimated dimensions of missing trees by averaging diameter, height and volume of trees in the plot. Given the negligible tree dimensions at the time of plantation (2008) volumes of trees in 2014 were used as a measure of tree productivity (i.e. volume increment: VI). Stand dendrometric characteristics are summarized in table 1.

**Table 1.** Mean (minimal-maximal) values of tree height, tree circumference and plot basal area of maritime pine (*Ppin*) and silver birch (*Bpen*) in monocultures (*mo*) and mixed stands (*mx*) at the three stand densities studied: low (625 t/ha), medium (1250 t/ha) and high (2500 t/ha).

|  |  | Stand density |  |  |  |  |
| --- | --- | --- | --- | --- | --- | --- |
|  |  | Low<br>Mo | Medium<br>Mo | Mx | High<br>Mo | Mx |
| Plot basal area (m<sup>2</sup>/ha) | *Bpen* | 1.41<br>0.84-2.25 | 2.39<br>0.80-4.62 |  | 3.55<br>1.71-5.10 |  |
|  | *Ppin* | 7.22<br>6.17-8.30 | 11.4<br>4.69-14.7 |  | 17.2<br>11.3-21.3 |  |
|  | *Bpen + Ppin* |  |  | 7.42<br>4.54-9.10 |  | 11.2<br>8.24-14.7 |
| Tree height (cm) | *Bpen* | 526<br>85-764 | 537<br>120-900 | 513<br>106-795 | 517<br>95-791 | 508<br>148-729 |
|  | *Ppin* | 570<br>232-706 | 575<br>133-801 | 581<br>250-780 | 571<br>200-810 | 567<br>161-795 |
| Tree circumference at<br>breast height (cm) | *Bpen* | 16.6<br>2.4-26.7 | 14.9<br>1.52-28.2 | 13.3<br>1.5-28.5 | 13.0<br>2.3-24.2 | 12.0<br>1.1-22.5 |
|  | *Ppin* | 37.6<br>9.5-51 | 33.4<br>0.7-50.2 | 36.1<br>10.1-49.9 | 29.1<br>5.3-45.2 | 30.6<br>2.6-47 |

### TRANSGRESSIVE MIXTURE EFFECT AND MIXTURE EFFECTS FOR SPECIES SUBSTITUTION AND ADDITION

We calculated two integrated indices of mixture effect for heterospecific addition and species substitution: mixture effect (ME) and transgressive mixture effect (TME). These indices were adapted from the classic calculation of transgressive overyielding and overyielding. Transgressive overyielding and overyielding are two standardized indices of mixture effect on stand productivity calculated by comparing monocultures to mixed stands at a same stand density (i.e. in case of species substitution). Yet, the major difference between species substitution and species addition is that total stand density increases from monoculture to mixed stands in an additive design, while it is kept constant in a substitutive design. It follows that the reference monoculture used to calculate ME and TME differed between additive and substitutive designs. We calculated TME the same way for species substitution and species addition at medium and high stand density (n = 2500 t/ha and n = 1250 t/ha) by comparing the mean total stand volume increment (SVI) of stands between mixed stands (*mx*) and monoculture stands (*mo*) of the most productive species, i.e. maritime pine (Fig. 2):

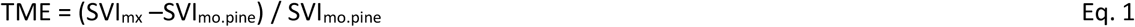

Where SVI_mx_ was the stand volume increment since plantation of mixed stands averaged per block; SVI_mo_ the stand volume increment in monoculture of birch or pine averaged per block.

**Figure 2.**
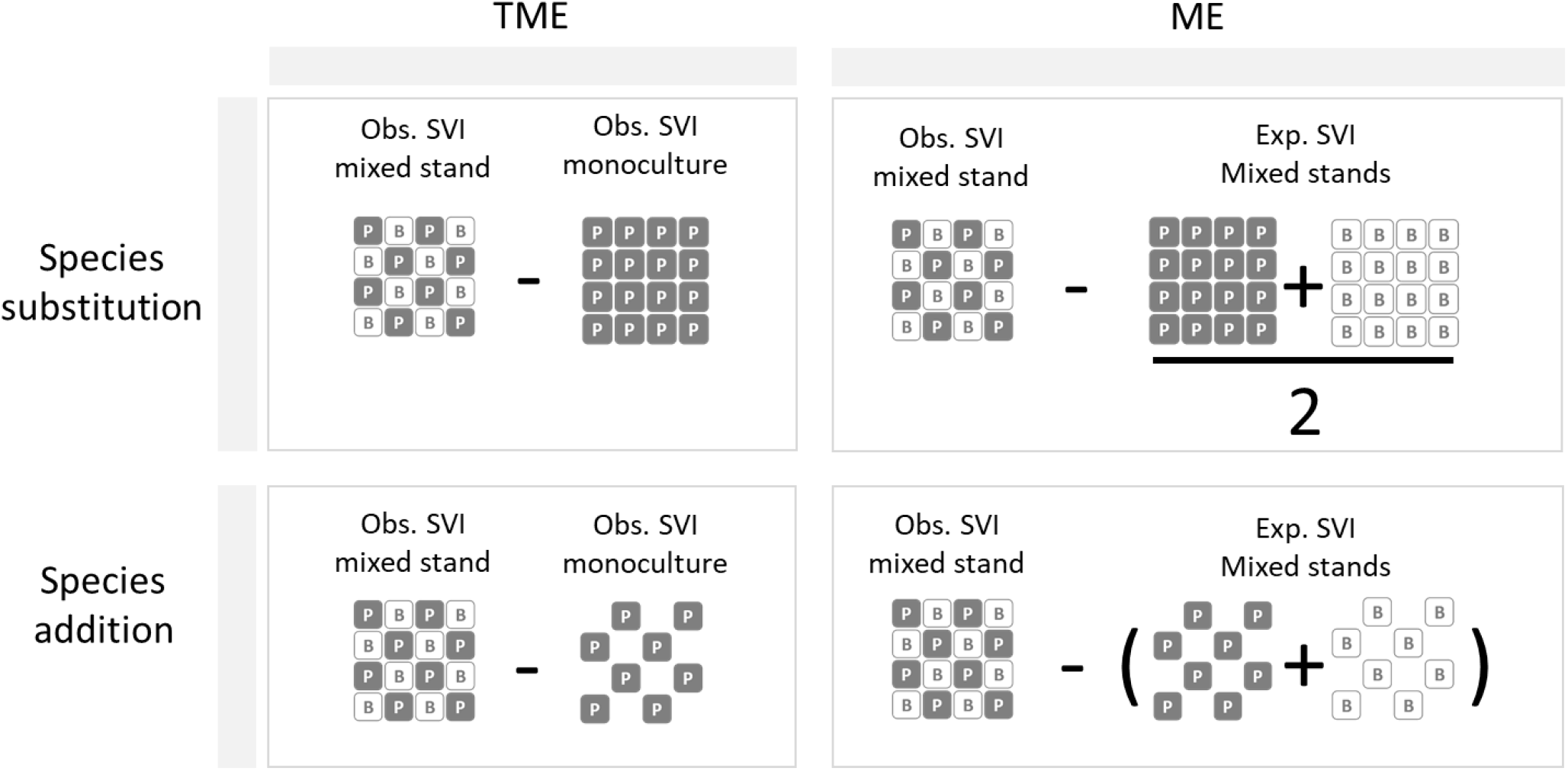
Schematic representation of the calculations of transgressive mixture effect (TME) and mixture effect (ME) of stand volume increment (SVI) for species substitution and species addition based on observed (obs.) values and expected (exp.) values.

The mixture effect (ME) was calculated for each block separately at medium and high levels of stand density (n = 2500 t/ha and n = 1250 t/ha) as:

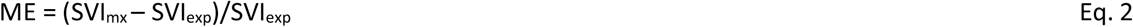

where SVI_mx_ was the observed volume increment of mixed pine-birch stands and SVI_exp_ was the expected volume increment of mixed stands. While SVI_mx_ was the same for both species substitution and species addition, SVI_exp_ differed between the additive and substitutive scenarios.

For species substitution, the calculation of SVI_exp.sub_ was:

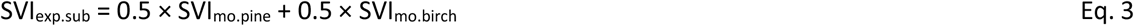

where SVI_mo. pine_ and SVI_mo. birch_ were the stand volume increment of pine and birch monocultures averaged per block and 0.5 corresponds to the species proportion.

For species addition, we compared SVI_mx_ to SVI_exp.add_ at an equal number of trees per species. Thus, for a SVI_mx_ at a density of n trees, we derived SVI_exp.add_ by summing volume increments in monocultures of n/2 trees (see Figure 2):

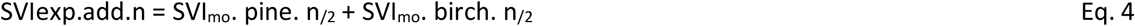

### STATISTICAL ANALYSES

All the analyses were performed with R 3.2.5 and functions gam, lme and glht in packages mgcv, nlme and multcomp.

We conducted separate analyses at the stand and tree levels by fitting a set of linear mixed effect models. At the plot level, we analysed four response variables: (i) total stand volume increment (SVI) estimated by summing tree volumes at plot level, (ii) mixture effect (ME) resulting from a species substitution (ME_sub_) and from a species addition (ME_add_) and (iv) transgressive mixture effect (TME) resulting from a species substitution (TME_sub_) and from a species addition (TME_add_). We completed the analyses at the plot level by considering also tree volume increment (TVI) of maritime pine and silver birch individual trees in monocultures and mixed plots.

Models of SVI and TVI included the effects of stand density (low, medium and high) and tree diversity (monoculture vs. two-species mixture) as fixed effect factors. Models of ME and TME included the effects of stand density (high and medium) and mixture scenario (substitution vs. addition) as fixed effect factors. We added Block as a random effect estimating between-block variability, except for analyses conducted at the level of individual tree where we nested plot within block to account for the non-independence of multiple trees sampled within the same plots and blocks.

To consider the residual heteroscedasticity, analyses of SVI and TVI were carried out by introducing a variance model into the linear mixed models allowing unequal variance among experimental treatments (Pinheiro & Bates, 2006).

## RESULTS

### TREE SPECIES SUBSTITUTION

The substitution of silver birch by maritime pine multiplied significantly SVI by 3.6 at medium stand density (Figure 3, 38.8±5.67 m^3^/ha, n=18; S1 in Figure 1) as well as at high stand density (Figure 3, 56.6±10.6 m^3^/ha, n=8; S2 in Figure 1). Conversely, species substitution of maritime pine by silver birch decreased significantly SVI by 35% (Figure 3, S3 in Figure 1) and 36% (Figure 3, S4 in Figure 1) at medium and high stand density.

**Figure 3.**
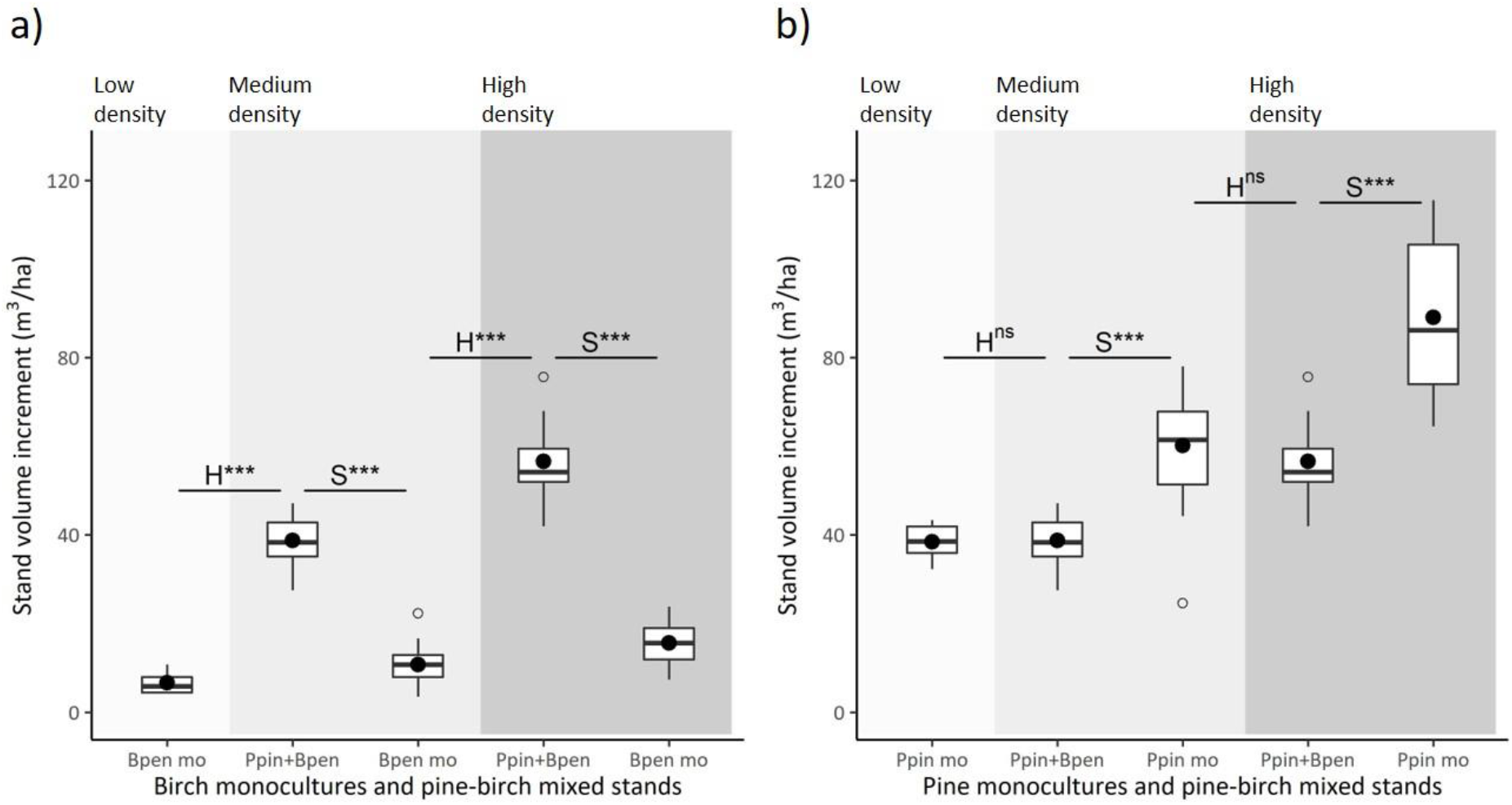
Stand volume increment (SVI) for different plot composition: in monoculture (mo) the SVI is the sum of tree volume increment of the target species; in mixed plots (Ppin + Bpen) the SVI cumulates the tree volume increment of silver birch (Bpen) and maritime pine (Ppin). Effect of heterospecific addition (H, see solid arrows in Figure 1) and species substitution (S, see black arrows in Figure 1) on stand volume increment (SVI) of silver birch (a) and maritime pine (b) at low, medium and high stand density. Black dots indicate mean values. Significance at a level of 5% were indicated by stars ((.) 0.1 > p-values > 0.05; * 0.05 > p-values > 0.01; ** 0.01 > p-values > 0.001; *** p-values > 0.001; ns no significant effect). Note that SVI P_pin + Bpen_ of the medium and high density mixed stands are the same in a) and b).

Birch-pine mixtures obtained through the substitution scenario were significantly less productive than the most productive (pine) monoculture (TME_sub_ < 0, Figure 4) at both medium (−0.35±0.08, n= 8) and high (−0.35±1.45, n=8) species density. Mixture effect (ME_sub_, figure 4) indicated that pine-birch mixtures was marginally significantly more productive (overyielding) than their component monocultures at medium (0.10±0.14, n=8) and at high stand density (0.10±0.20, n=8, Figure 4).

**Figure 4.**
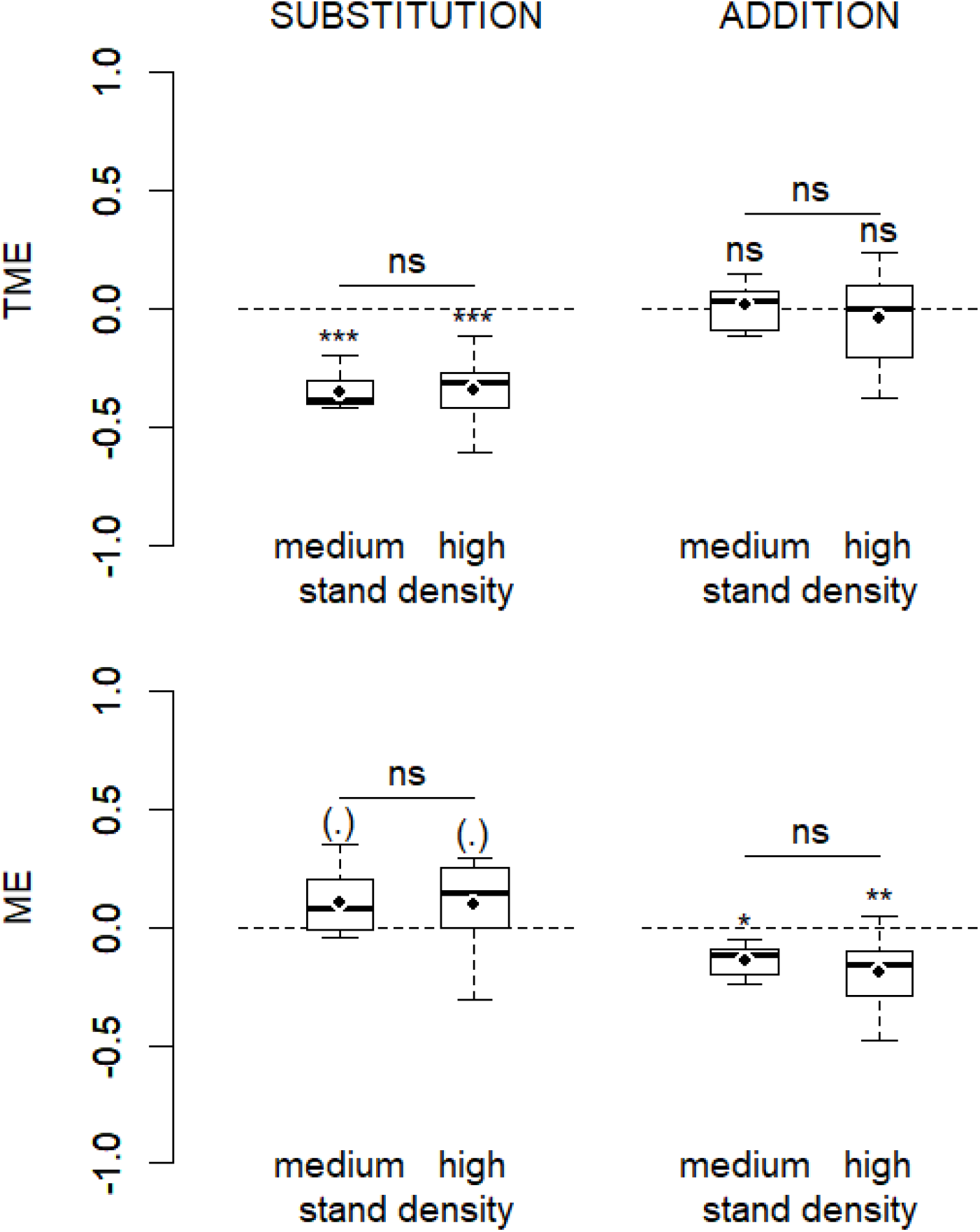
Transgressive mixture effect (TME) and mixture effect (ME) at the stand level for species substitution and species addition at medium and high stand density. Black dot indicates the mean effects. Significance at a level of 5% were indicated by stars ((.) 0.1 > p-values > 0.05; * 0.05 > p-values > 0.01; ** 0.01 > p-values > 0.001; *** p-values > 0.001; ns no significant effect)

Species substitution had opposite effects on TVI (tree volume increment) of the two studied species at medium stand density: a substitution of maritime pine by silver birch caused a significant increase of 15% in pine TVI (Figure 5), but a significant reduction of 23% in birch TVI (Figure 5). At high stand density, species substitution did not have any significant effect on TVI of silver birch or maritime pine (Figure 5).

**Figure 5.**
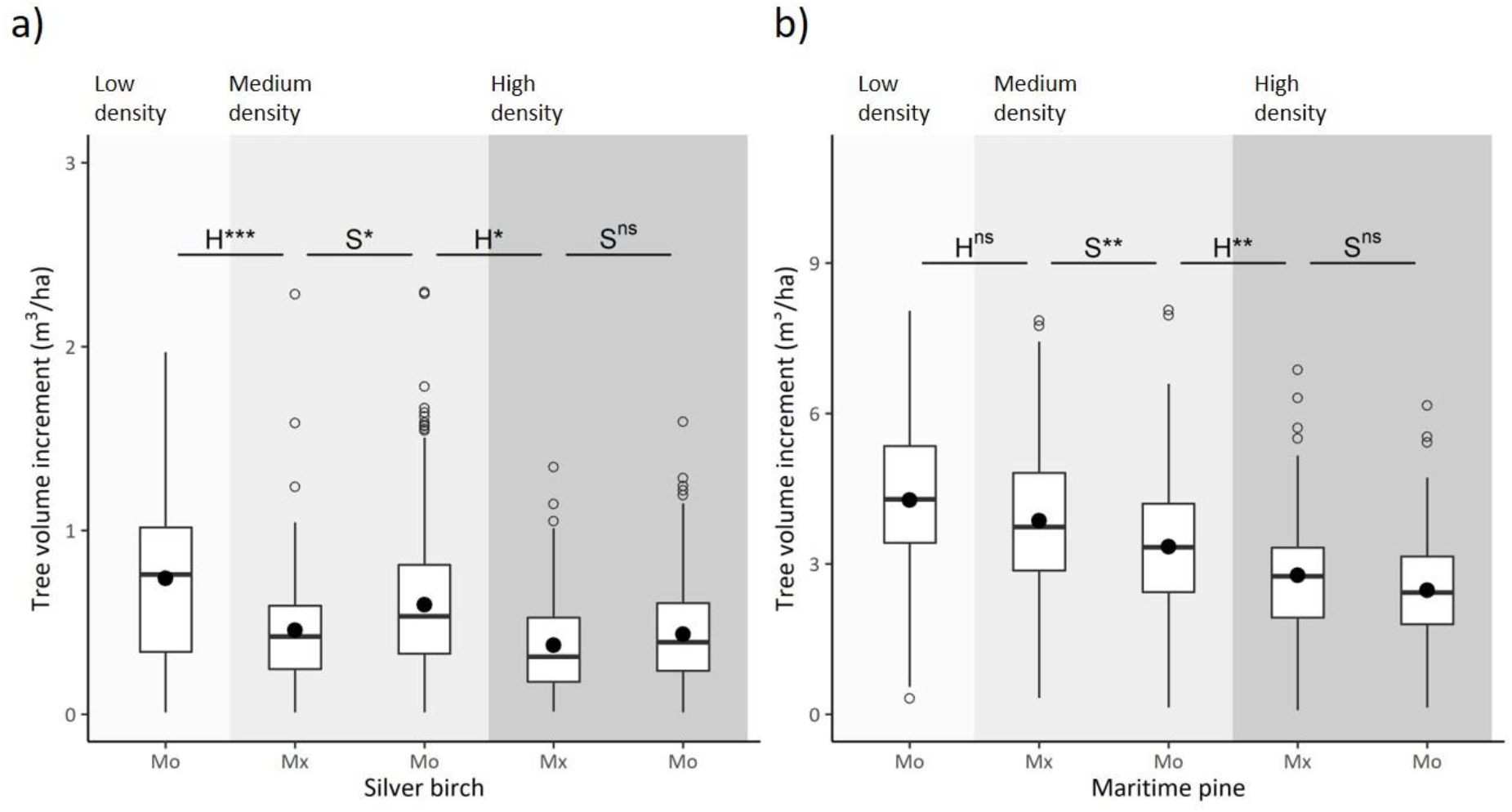
Effect of heterospecific species addition (H) and species substitution (S) on tree volume increment (TVI) of silver birch (*Betula pendula*) (a) and maritime pine (*Pinus pinaster*) (b) at low, medium and high stand density and in monoculture (Mo) or mixed (Mx) plots. Black dots indicate mean values. Significance at a level of 5% were indicated by stars ((.) 0.1 > p-values > 0.05; * 0.05 > p-values > 0.01; ** 0.01 > p-values > 0.001; *** p-values > 0.001; ns no significant effect). Note that the scales of the two figures (m^3^/ha) are different.

### HETEROSPECIFIC TREE ADDITION

Heterospecific species addition of maritime pine in silver birch stands multiplied significantly SVI by 5.8 at medium stand density (Figure 3, H1 in Figure 1) and by 5.2 at high stand density (H3 in Figure 1). Heterospecific species addition of silver birch in maritime pine stands did not have any significant effect on SVI neither at medium stand density (Figure 3, H3 in Figure 1) nor at high stand density (H4 in Figure 1).

ME_add_ indicated that pine-birch mixtures were significantly less productive (underyielding) than their component monocultures at intermediate (−0.14±0.07, n=8) and at high stand density (−0.19±0.16, n=8, Figure 4). TME_add_ at medium (0.01±0.10, n=6) and high (−0.05±0.20, n=8) stand density were not significantly different from zero indicating that SVI of mixed stands did not differ from SVI of pine in monoculture, i.e. no transgressive overyielding (Figure 4).

Heterospecific addition of silver birch in monoculture of maritime pine did not cause any significant change in TVI of maritime pine at medium stand density (Figure 5) but a significant reduction of 17% (Figure 5) at high stand density. Heterospecific addition of maritime pine in in monoculture of silver birch caused a significant reduction in TVI of silver birch of 42% (Figure 5) and 36% (Figure 5) at medium and high stand density, respectively.

## DISCUSSION

Our study assessed the role of tree species addition and species substitution in mixture effect in stands at an early age. We highlighted that when controlling for stand density, overyielding in young silver birch – maritime pine stands was due to a relaxation of intra-individual competition for pine. Conversely addition of silver birch (the least productive species) in a maritime pine stand (the most productive species) did not have a negative impact on stand productivity, which implies a non-significant transgressive mixture effect. Eventually stand density had little impact on the mixture effects tested and rather contribute to the species responses at an individual scale.

### 1§ SPECIES SUBSTITUTION TRIGGED OVERYIELDING IN MIXTURE OF TWO PIONEER SPECIES

Respective growth rates of tree species are crucial for interactions between species in the early stages of development of mixed forests and our results confirm that positive effects of biodiversity on productivity are mainly due to selection effect (Tobner et al., 2016) i.e. a fast-growth and productive species driving ecosystem functioning. Competitive advantage is common in young forests and positive diversity-productivity relationship at this stage are often attributed in a lesser extent to complementarity, particularly in harsher conditions (Van de Peer et al., 2018). Such positive effects are commonly attributed to differences in shade tolerance as species with rapid growth rates benefit from a relaxation of intraspecific competition, which may or may not be accompanied by niche separation favouring shade-tolerant species rapidly overtopped due to their lower height growth rate (Boyden et al., 2009; Tobner et al., 2016). However, we evidenced that overyielding can be triggered by species similarities in their shade tolerance. Mixture effect was not conditioned by different light acquisition strategies but more probably by their unequal ability to both tolerate drought. The experimental plantation was on sandy heathlands that experience intense drought episodes in summer, water availability is an important limiting factor for tree growth, especially in silver birch, which has the lowest drought tolerance. Maritime pine is able to maintain its stem growth during a longer period and even to restart height grow in autumn (fast-growth evergreen species, (Heuret et al., 2006)). Silver birch remains a species sensitive to interspecific competition at a young age, even in the Nordic countries where temperature is a more limiting factor for growth than water (Jucker et al., 2020), but it is likely that dry conditions further accentuate its competitive disadvantage.

Nonetheless effects that we observed 8 (7?) years after planting will change very quickly, the growing gap in height between maritime pine and silver birch being detrimental to birch under current climatic conditions, tree mortality will intensity (Morin et al., 2020). Long term simulations of pine and birch stand showed a lasting overyielding due to the relaxation of intraspecific competition for pine over time (Morin et al., 2020). Oaks species with their slower growth rates and varying drought and shade tolerances will gradually establish into stands, leading to a stand stratification possibly suitable to mixed stands, even though at this point, there is no consensus on the stage of forest development for which the positive effect of diversity peak (Jucker et al., 2020; Taylor et al., 2020).

### 2§ TRANSGRESSIVE MIXTURE EFFECT WAS NOT SIGNIFICANTLY DIFFERENT FROM ZERO IN THE ADDITION SCENARIO

We did not find any transgressive mixture effect in mixed birch-pine stand created by addition of the two species. Conversely the substitutive approach caused a loss of mixed stand productivity compares to pine monocultures due to the substitution of a high productive species (maritime pine) by a low productive species (silver birch). These findings mirror what has been found in colder and more humid sites for the same species that we studied (Frivold & Frank, 2002), more generally in mixed-forest (Jactel et al., 2018) and in plant community where positive transgressive overyielding were rarely reported (Cardinale et al., 2007).

At a medium stand density in the additive scenario the absence of any competition effect of silver birch on maritime pine can be explained by two inseparable mechanisms: either, a purely neutral effect of the addition of the least productive species due to an insufficient proximity of stems, or a facilitating effect of birch on soil resource that compensates for a weak competitive constraint due to species addition. Leaves of silver birch have a rate of decomposition higher than needles of Pinus species (Palviainen et al., 2004) and depending on the stand structure, nutrient cycling can be higher in birch regeneration than in pine regeneration (De Schrijver et al., 2009). Hence in the studied site, carbon and nitrogen at an intermediate soil depth have been found higher in mixed stands than on monospecific stands (Maxwell et al., 2020), even though there is no evidence on belowground complementarity of fine roots (Altinalmazis-Kondylis et al., 2020).

These findings are also of great ecological relevance because they demonstrate that it is possible to diversify pine monocultures with addition of birch at an early age and then benefit from ecosystem services such as pest protection (Damien et al., 2016; Jactel et al., 2019) and increased diversity of predatory insects (Jouveau et al., 2020) without compromising the wood production of the target species (here maritime pine). Long-term simulations of pine and birch growth on the study sites supported our results (Morin et al., 2020) showing that the ecosystem services associated with pine monoculture diversification can persist as the stand ages.

### 3§ MIXTURE EFFECTS AND TRANSGRESSIVE MIXTURE EFFECTS DO NOT CHANGE WITH STAND DENSITY, BUT TREE PRODUCTIVITY DOES

In young stands, high stand densities usually speed up mixture effects (Tobner et al., 2016; Van de Peer et al., 2018). In this study, we did not observe any intensification with stand density neither of the mixture effect nor of transgressive mixture effect. However, an intensification of interactions with stands density was observed at the tree level: at medium density, heterospecific addition did not affect maritime pine trees (the most productive species) but silver birch trees (the least productive species). At high density, the intensification of interspecific competition led to a reduction of productivity for both species. Regarding species substitution, at medium density maritime pine (the most productive species) benefit from mixture at the expense of silver birch (the least productive species). When stand density increased, the difference between inter and intraspecific competition becomes narrower, which might explain the absence of any effect of species substitution. This illustrates on the one hand that the same response pattern in term of mixture effect can emerge from different mechanisms at the individual levels and. On the other hand, it is consistent with what has been observed elsewhere in young and dynamic stages with an intensification of competitive interactions (Boyden et al., 2009) or at least a decrease in overyielding with density (Kweon & Comeau, 2019). Finally, these results contrast with the intensification of the positive diversity productivity relationship observed as forest stands become older (Huang et al., 2018) particularly when shade tolerant and shade intolerant species are mixed (Brunner, 2020; del Rio & Sterba, 2009).

## CONCLUSION

By controlling stand density and species identity, we have shown that selection effect is the main driver of positive diversity – productivity relationships in the early stages of mixed species forests. This calls for a careful choice of which tree species to associate when designing plantations of mixed species, especially of fast-growing species. In addition, our results also showed that the addition of a pioneer and low-demand species to a monoculture of a high productive species in young developmental stages offers the opportunity to benefit from ecosystem services associated to mixed stands without affecting the productivity of the target species. The addition of tree species is a promising way to promote multifunctionality in mixed-species plantations and to preserve the harvest of a particular species for timber production, thus circumventing two major obstacles in the implementation of mixed-species forestry.

## ACKNOWLEDGEMENTS

We thank the Forest experimental Facility (UEFP-https://doi.org/10.15454/1.5483264699193726E12) and especially Bernard Issenhuth for the maintenance and the measurements of the ORPHEE experiment

## AUTHORS’ CONTRIBUTIONS

All authors contributed critically to conceived the ideas and designed methodology; HJ designed the ORPHEE experiment; MT analysed the data; MT and CM led the writing of the manuscript. All authors contributed critically to the drafts and gave final approval for publication.

## DATA AVAILABILITY STATEMENT

Data will be available from the INRAE Digital Repository https://data.inrae.fr/dataverse/biogeco.

