## supplementary material for "Tree diversity effects on forest productivity: disentangling the effects of tree species addition vs. substitution"

Supporting Information Table S1. Number of plots per treatment for different threshold of tree mortality.

| | | | Percentage of dead trees | | | | | | |
|---|---|---|---|---|---|---|---|---|---|
| Stand density | Stand composition | Total number of plots | 30% | 25% | 20% | 15% | 10% | 5% | 1% |
| 2500 t/ha | Bpen | 8 | 8 | 8 | 8 | 8 | 8 | 7 | 3 |
| | Bpen+Ppin | 8 | 8 | 8 | 8 | 8 | 8 | 8 | 8 |
| | Ppin | 8 | 8 | 8 | 8 | 8 | 8 | 7 | 5 |
| 1250 t/ha | Bpen | 24 | 24 | 24 | 23 | 23 | 23 | 18 | 18 |
| | Bpen+Ppin | 24 | 24 | 24 | 23 | 23 | 20 | 20 | 20 |
| | Ppin | 24 | 24 | 24 | 23 | 23 | 22 | 18 | 18 |
| 625 t/ha | Bpen | 8 | 8 | 8 | 8 | 8 | 6 | 6 | 6 |
| | Ppin | 8 | 7 | 7 | 6 | 6 | 5 | 5 | 5 |

Supporting Information Figure S1. Estimation of the circumferences of trees.

Height-circumference relationships are known to change with tree age, stand composition and tree density, we thus fitted a separate generalized additive model for each stand composition, density and year (Appendix 1). Relationships between tree circumference (cm) and tree height (cm) in the 8 blocks of the experimental design. The grey dots represent the raw data and lines represent the predicted values for *Pinus pinaster* and *Betula pendula* at 650 t/ha, 1250t/ha and 2500 t/ha in monocultures and mixed stands.

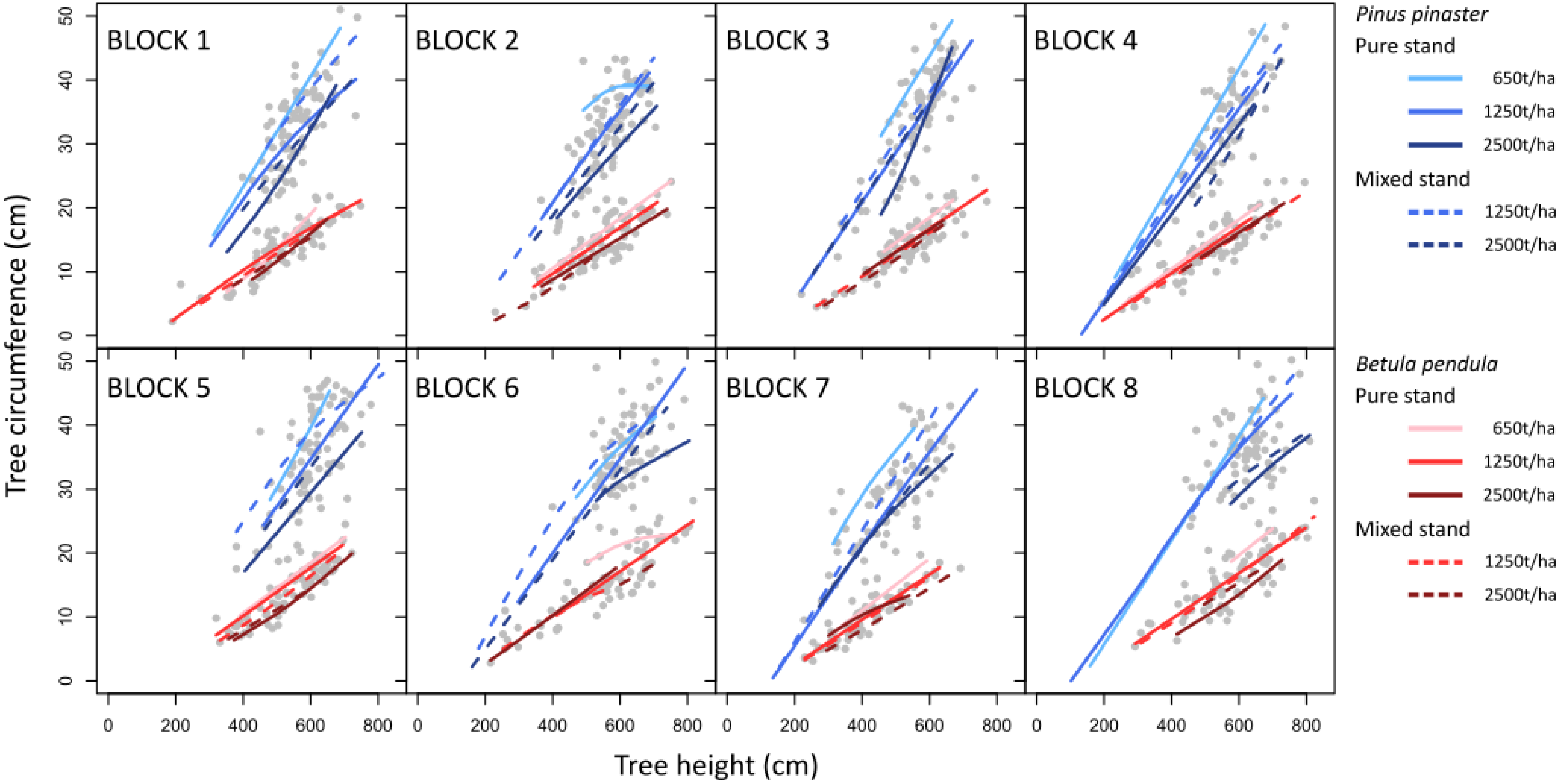
